# Lung lipid deposition in pneumonias of different etiologies

**DOI:** 10.1101/2022.12.30.522299

**Authors:** Daria M Potashnikova, Anna V Tvorogova, Alexey A Komissarov, Aleena A Saidova, Tatiana N Sotnikova, Valeria O Makarova, Eugene A Arifulin, Tatiana V Lipina, Olesya M Shirokova, Eugene S Melnikov, Tatiana A Rodina, Anna A Valyaeva, Anastasia A Zharikova, George O Zayratyants, Oleg V Zayratyants, Eugene V Sheval, Leonid B Margolis, Elena J Vasilieva

**Affiliations:** City Clinical Hospital Named After I.V. Davydovsky, Moscow Department of Healthcare, Moscow, Russia; A.I. Yevdokimov Moscow State University of Medicine and Dentistry of the Ministry of Healthcare of the Russian Federation, Moscow, Russia; Department of Cell Biology and Histology, School of Biology, Lomonosov Moscow State University, Moscow, Russia; Belozersky Institute of Physico-Chemical Biology, Lomonosov Moscow State University, Moscow, Russia; Central Scientific Research Laboratory, Institute of Fundamental Medicine, Privolzhsky Research Medical University, Nizhny Novgorod, Russia; Department of Bioengineering and Bioinformatics, Lomonosov Moscow State University, Moscow, Russia; City Clinical Hospital Named After S.S. Yudin, Moscow Department of Healthcare, Moscow, Russia; Faculty of Natural Sciences and Medicine, Ilia State University, Tbilisi, Georgia

**Author notes:** Corresponding authors: E.J. Vasilieva, D.M. Potashnikova.

**Keywords:** lung pathology, pneumonia, COVID-19, lipid deposition, lipid metabolism

## Abstract

Pneumonia is an acute respiratory disease of varying etiology that has drawn much attention during the COVID-19 pandemic. Among the many thoroughly studied aspects of pneumonia, lipid metabolism has not been sufficiently addressed. Here, we investigated lipid deposition in the *post mortem* lung specimens of patients who died from COVID-19 and non-COVID-19 pneumonias. We used semi-thin sections and cryosections stained with Sudan III to visualize lipid droplet deposition within cells and in the extracellular space, most notably in small lung vessels. Electron microscopy analysis of the ultrathin sections was used to confirm the homogeneous structure of the droplets. Morphometric analysis revealed that the area of lipid deposition was increased in pneumonia compared to control lung tissue. Likewise, it was increased in the macroscopically inflamed vs. the macroscopically intact area of the same pneumonia lung. The lipid profiling by chromato-mass spectrometry revealed that lipid droplet accumulation in pneumonia was associated with a specific fatty acid content of the inflamed lung tissue. The gene expression analysis pointed to changes of lipid metabolism in the inflamed lung tissue compared to control lungs. Taken together, our data indicate a number of morphologic and metabolic changes associated with inflammation and common for pneumonias of different etiologies that likely contribute to pneumonia pathogenesis. Therefore, targeting lipid metabolism can be considered a new therapeutic strategy.

## Introduction

Pneumonias of various etiologies are life-threatening conditions that affect alveoli and distal bronchioles, or interstitial lung tissue. Many morphological [1,2] and physiological [3,4] abnormalities have been reported in inflamed lungs. However, dysregulation of lipid metabolism associated with lung inflammation has been addressed much less, although lipid synthesis in general and lung surfactant specifically have an important protective role in lung diseases [5]. Recently, different aspects of pneumonia have attracted attention in the context of COVID-19, as the lungs are one of the main target organs of SARS-CoV-2 [6]. However, the lipid metabolism in the context of this infection has not been thoroughly studied.

Here, we addressed this issue by investigating lipids in postmortem lung specimens from individuals who died of pneumonia. The lipids in inflamed lung tissue were deposited as droplets in the cell cytoplasm, in the extracellular space, and within small blood vessels. These lipid depositions were largely restricted to the inflamed area of the lung and were associated with elevated levels of unsaturated fatty acids and generally decreased expression of genes involved in lung lipid metabolism.

To determine whether these lipid abnormalities are characteristic of COVID-19–associated pneumonia, we extended our study to pneumonias of different etiologies. In the lungs of SARS-CoV-2–negative patients who died of pneumonia, we found lipid depositions similar to those in SARS-CoV-2–positive patients. Abnormal lipid deposition may contribute significantly to the development of pathological processes associated with pneumonia.

## Materials and methods

### Autopsies and post-mortem sampling

The first part of the study included lung autopsies from 12 patients who died of COVID-19-associated pneumonia. The analysis included qPCR verification of SARS-CoV-2 in *post mortem* tissue, H&E section assessment, semi-thin and ultrathin specimen assessment.

The second part of the study included *in silico* analysis of the available RNA-seq databases using Gene Ontology to highlight the metabolic pathways altered in the COVID-19-associated pneumonia.

The third part of the study included the examination of lung autopsies from 26 (16 pneumonia and 10 control) individuals. The analysis included macroscopic assessment of *post mortem* tissue, qPCR verification of SARS-CoV-2 in pneumonia, H&E section assessment, Sudan III staining assessment, qPCR expression analysis of genes involved in lipid metabolism.

The fourth part of the study included examination of the paired lung specimens (from the area with macroscopically distinct inflammation in the inferior lobe of the lung (segments S8 or S9), and from the macroscopically intact area in the superior lobe of the lung (segments S1 or S2)) obtained from COVID-19-associcated pneumonia autopsies and non-COVID-19-associated pneumonia autopsies. The non-pneumonia lungs served as negative control. The analysis included macroscopic assessment of *post mortem* tissue, qPCR verification of SARS-CoV-2 in pneumonia, H&E section assessment, Sudan III staining assessment and morphometry as well as chromato-mass spectrometry of fatty acids in the specimens.

The study was approved by the Moscow City Ethics Committee (protocol № 50/69_13.10.2020) of the Research Institute of the Organization of Health and Healthcare Management and performed according to the Declaration of Helsinki. All specimens were obtained from the City Clinical Hospital named after I.V. Davydovsky and the Department of Pathology of the City Clinical Hospital named after S.S. Yudin re-profiled for COVID-19 autopsies according to the international and Russian requirements for working with COVID-19. The autopsy data are provided in Supplementary Table S1.

### Histological specimens

All dyes for histological staining were provided by Labiko (Moscow, Russia). Lung autopsy specimens 1.0×1.5×0.5 cm were fixed in 10% buffered formalin. For routine hematoxylin and eosin (H&E) analysis the specimens were dehydrated and cast in paraffin blocks according to the standard procedure. Paraffin sections 3-4 μm thick were stained with H&E, dehydrated and mounted in Vitrogel (Biovitrum, Russia).

For Sudan III staining 7 μm thick cryosections of formalin-fixed tissue were made using a Leica CM1850 cryotome (Leica, Germany), placed on Superfrost Plus glass slides (Thermo Scientific, US) stained with Sudan III by Herxheimer according to the manufacturer’s protocol, co-stained with Carazzi hematoxylin (exposure time 40 sec, three rinses in tap water) and mounted in glycerol.

Specimens were imaged using a Leica DM2000 microscope with Leica DFC7000 T camera (Leica, Germany).

High-power frames were made with x40 objective.

### Pathological analysis on semi- and ultrathin sections

Lung autopsy specimens 1×1×1 mm in size were fixed in 2.5% glutaraldehyde (Ted Pella, US) in cacodylate buffer, post-fixed in 1% OsO_4_, dehydrated in ethanol and propylene oxide (Sigma-Aldrich, US), and embedded in Epon (Fluka, Switzerland). We obtained semi-thin sections (0.5 µm thick) with a Leica ultramicrotome using glass knives and stained them on glass slides using methylene blue/azure II or polychromatic Twort’s staining, as described earlier [7] (Supplementary Figure S1A–B). The sections were mounted in Epon resin under a coverslip, and imaged using a Leica DM2000 microscope equipped with a Plan-Neofluar 100/1.3 Oil immersion objective and a Leica DFC7000 T camera.

Ultrathin sections were made using a diamond knife and mounted on slot grids (Ted Pella, US), stained with uranyl acetate and lead nitrate, and examined using a JEM-1400 electron microscope (Jeol, Japan) at an accelerating voltage of 80 kV.

### Immunohistochemistry specimens

Cryosections of formalin-fixed lung tissue 7 μm thick were washed in PBS and permeabilized for 1 h with 0.01% Tween 20 (Sigma-Merck, Germany), followed by three washes in PBS. We stained the specimens for vascular endothelium using a CD31-FITC antibody (clone WM59, BD Biosciences, US) at 1:100 dilution in a humid chamber at +37ºC for 1 h. Oil Red O (Sigma-Merck, Germany) dye was used for staining lipids; sections were stained for 20 min at RT according to the manufacturer’s protocol. Nuclei were co-stained with DAPI (Thermo Scientific, US) at a final concentration of 1μg/ml. Fluorescence imaging of ORO/anti-CD31/DAPI staining was performed in scanning mode using an inverted Zeiss AxioObserver (Zeiss, Germany) microscope operating under Zen 3.1 Blue Edition software with a ×40/Oil objective, standard filter cubes and Hamamatsu ORCA-Flash4.0 V2 camera (Hamamatsu Photonics, Japan). We used ImageJ software (NIH) for image processing.

### Morphologic and morphometric analysis of Sudan III-stained specimens

Morphologic analysis was performed on H&E sections and included a blinded examination of the images by three independent pathologists. The absence or presence of alveolar damage characteristic of pneumonia were confirmed for non-pneumonia and pneumonia autopsies respectively.

The initial morphologic analysis of lipid depositions was performed on Sudan III-stained cryosections counterstained with hematoxylin. The further morphometric comparison of inflamed and intact pneumonia lung tissue and non-pneumonia lung tissue was performed on selected Sudan III-stained cryosections at 20 high-power fields per each specimen imaged with objective x40. Image Pro software (Media Cybernetics, US) was used. For each image, lipid droplets were included in a mask containing pixels by color, adjusted to include all Sudan III–positive structures into the regions of interest (ROIs). The resulting ROIs were assessed as area per image.

### Chromato-mass spectrometry

To determine the fatty acid (FA) profile, we analyzed 10 pneumonia lung specimens. A tissue section was taken from the area with macroscopically distinct inflammation and from the macroscopically intact area of the same lung. Lung tissue was finely cut, aliquoted, weighed, and stored at −40°C prior to analysis. One aliquot was used to determine the moisture content of the sample as the weight loss on lyophilization, another aliquot was used to determine the FA profile. Thawed specimens were homogenized using glass beads. We extracted lipids using the Folch method [8] and determined the content of 31 FAs using a gas chromatograph mass spectrometer Shimadzu TQ8050 (Shimadzu, Japan). We used chemical ionization with ammonia [9], and chromatographic separation using a DB-23 column (60 m, 0.25 mm, 0.15μm, 122-2361 Agilent, US). FAs were identified on the basis of the retention times of analytes and the fragmentation reactions of quasimolecular ions of carboxylic acid ethyl esters and/or their adducts with ammonia [9]. The total mass of FAs in the specimen and their mass fractions were calculated.

### RNA extraction and reverse transcription

For total RNA extraction, one lung section 5×5×5 mm in size was finely cut and stored at −20°C in RNAlater (Thermo Scientific, US). We mechanically disrupted the tissue from RNAlater using a MediMachine (BD Biosciences, US) in 700 μL of RLT buffer (Qiagen, Germany) with 1% β-mercaptoethanol (Merck, Germany). The volume of tissue suspension was adjusted to 1.5 mL with nuclease-free water. The homogenate was incubated with 200 μg/mL Proteinase K (Thermo Scientific, US) at +54°C for 50 min and centrifuged for 10 min at 10,000 g. The supernatant was processed according to the RNeasy mini kit protocol (Qiagen, Germany). We eliminated DNA by on-column digestion using RNase-free DNase (Qiagen, Germany). We reverse-transcribed cDNA from total RNA using the ImProm II Kit (Promega, US) with Random primers, according to the manufacturer’s instructions.

### Real-time qPCR

We detected SARS-CoV-2 RNA in total RNA specimens using TaqMan one-step qPCR. We analyzed amplification of two different regions of the SARS-CoV-2 nucleocapsid gene (*N2* and *N3*), simultaneously using primers described earlier [10]. Additionally, the ubiquitin C (*UBC*) gene was used as an internal control in a triplex reaction. The primers and probes are listed in Supplementary Table S2. RNA solution (5 μl per well) was mixed with 5 μl of triplex primer/probe mix and 10 μl of qScript XLT One-Step RT-qPCR Tough Mix (Quantabio, US).

The genes involved in lipid storage and metabolism (*LPL, PLA2G4A, PLA2G2A, PLIN1, PLIN2, ACSL1, ACSL3, ACSL4, ACSL5, ACSL6*), were analyzed in cDNA specimens using a Syntol PCR kit with SYBR Green I (Russia). The primers for the genes are listed in Supplementary Table S3. The relative quantities of target genes were normalized to the ubiquitin C (*UBC*) reference gene. We analyzed all samples in triplicate using a CFX96 Touch cycler with CFX-Manager software (Bio-Rad, US).

### Meta-analysis of bulk and single-cell RNA-seq datasets

Published bulk and single-cell RNA-seq data from human COVID-19 lung tissue were re-analyzed to identify lipid metabolism genes with altered expression in pneumonia samples compared to healthy controls. The bulk RNA-seq dataset GSE150316 [11] was obtained from the Gene Expression Omnibus (GEO) database. Single-cell RNA-seq data were downloaded from the SCovid database [12], which accumulated datasets from different studies as lists of differentially expressed genes. We used clusterProfiler (version v3.18.1) [13] to perform functional over-representation analysis and focused on Gene Ontology (GO) terms enriched with differentially expressed genes related to lipid and fatty acid metabolism. Briefly, we used keywords “lipid,” “fat,” “triglyceride,” or “cholesterol” to search the enrichment results for the specific GO categories. For single-cell RNA-seq data, the REVIGO tool [14] was used to remove redundant GO terms.

### Statistics

The data were plotted and analysed using the R programming language (R Core Team (2023). R: A Language and Environment for Statistical Computing. R Foundation for Statistical Computing, Vienna, Austria. https://www.R-project.org/), RStudio software (Posit team (2023). RStudio: Integrated Development Environment for R. Posit Software, PBC, Boston, MA. URL http://www.posit.co/), and GraphPad Prism 8 software.

## Results

### Lipid droplet deposition is associated with COVID-19 pneumonia

We examined 12 H&E sections of COVID-19 lung autopsies. The diagnosis of COVID-19 was established by qPCR of SARS-CoV-2 RNA before/during hospitalization of the patients and later confirmed by us by qPCR on post-mortem lung tissue. Histological examination revealed all typical changes associated with pneumonia alveolar damage (Supplementary Figure S1 A-E).

Apart from the multiple morphological alterations characteristic of pneumonia round cavities surrounded by erythrocytes were often observed in congested small lung vessels (Figure 1A-B): a diligent search revealed multiple examples of such cavities, even in a single H&E specimen (Supplementary Figure S2). We hypothesized that they might contain lipid droplets that were removed during routine ethanol dehydration.

**Figure 1.**
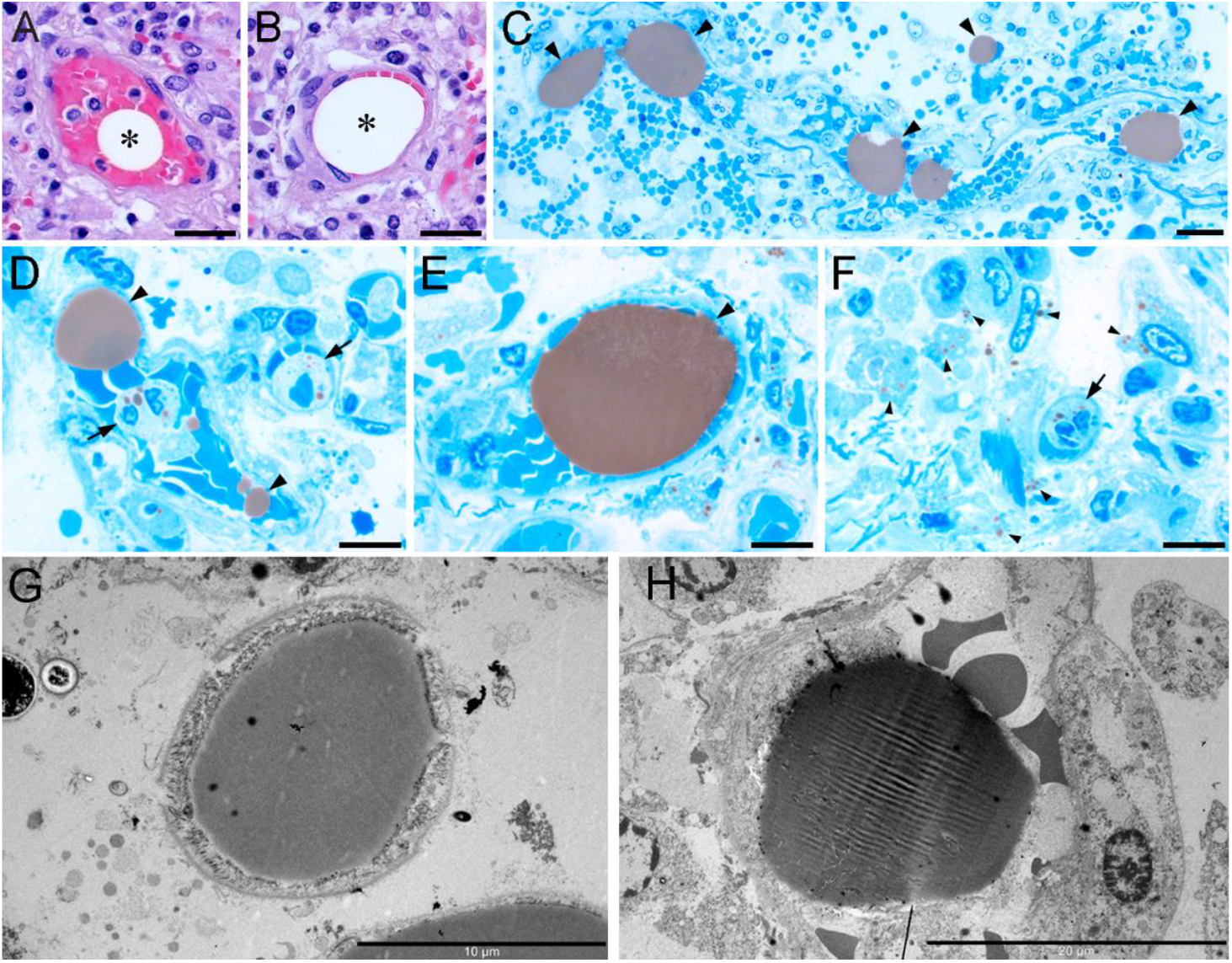
Lipid droplets in COVID-19–associated pneumonia lung. A. Paraffin section, hematoxylin and eosin staining. A small vessel contains a round cavity surrounded by erythrocytes. Scale bar, 25 μm. B. Paraffin section, hematoxylin and eosin staining. Erythrocytes in a small vessel are deformed and pushed to the vessel wall. Scale bar, 25 μm. C. OsO4-contrasted lipid droplets inside blood vessels in the lung. Scale bar, 100 μm. D. OsO4-contrasted lipid droplets are present in the blood vessel (arrows) and in the cells near the vessel (arrowheads). Scale bar, 10 μm. E. OsO4-contrasted lipid droplet fills the blood vessel (arrow), with erythrocytes pushed to the vessel wall. Cells and cell remnants in the vicinity of the vessel contain smaller lipid droplets (arrowheads). Scale bar, 10 μm. F. OsO4-contrasted small lipid droplets in the cytoplasm of the lung-infiltrating neutrophil (large arrowhead) and in other intact and destroyed cells (small arrowheads). Scale bar, 10 μm. G. OsO4-contrasted lipid droplets are homogeneous in ultrathin sections. EM, Scale bar, 10 μm. H. A homogeneous OsO4-contrasted lipid droplet in the blood vessel with erythrocytes; the erythrocytes are pushed to the vessel wall. EM, Scale bar, 20 μm.

To confirm this, we analyzed semi-thin lung sections post-fixed with OsO_4_. Two staining methods of the semi-thin sections were compared: methylene blue/azure II and Twort’s polychromatic staining. As the OsO_4_-contrasted lipid droplets appear close in color to erythrocytes upon Twort’s polychromatic staining (Supplementary Figure S3) we decided to further use methylene blue/azure II where the lipid droplets are best contrasted against other tissue elements. Examination of 12 semi-thin specimens of lung tissue clearly revealed large (10–100μm in diameter) OsO_4_-contrasted lipid droplets inside blood vessels (Figure 1C-E). These droplets either filled the entire volume of the vessel (Figure 1C) or were located among erythrocytes (Figure 1D-E), which corresponded to our findings in the paraffin sections. We reliably identified droplets in blood vessels in 6 out of 12 autopsy specimens, which mostly corresponded to late stages of the pathological process in the lungs. Additionally, we found numerous cells with small lipid droplets in the cytoplasm. Some cells (neutrophils by morphology) were located inside the blood vessels and some (macrophages by morphology) – inside the lung tissue (Figure 1F).

Electron microscopy (EM) of lipid droplets inside the blood vessels revealed their homogeneous structure (Figure 1G–H). The erythrocytes in contact with these droplets changed their shape, indicating significant mechanical toughness and stability of these droplets.

### Lipid metabolism in the lung is affected in COVID-19 pneumonia

To investigate the metabolic processes underlying lipid accumulation in lung tissue, we employed Gene Ontology (GO) analysis of the publicly available bulk and single-cell RNAseq datasets. The bulk RNAseq analysis of nine independent COVID-19 cases and the cumulative dataset revealed the overall trend for gene downregulation in infected lung, despite the high inter-specimen variability (Supplementary Figure S4). All of the specimens had alterations in the pathways concerning lipid metabolism, most commonly in the pathways of “fat cell differentiation” (downregulated in 6 specimens) and “regulation of lipid metabolic process” (downregulated in 5 specimens). The cumulative dataset retained but two downregulated pathways, namely “regulation of lipid biosynthetic process” and “positive regulation of lipid biosynthetic process”.

At single-cell level, the tendency was confirmed: the downregulated genes prevailed in lung cells, most notably in fibroblasts, macrophages, and T cells (Supplementary Table S4). Among the downregulated genes in macrophages and endothelial cells, we found ones involved in free fatty acid / cholesterol uptake, storage, and metabolism. Together these data suggest that COVID-19 pneumonias are associated with shifts in lipid metabolism and the accumulation of lipid droplets in cell cytoplasm and in the small blood vessels of the lung.

### Quantification of lipid depositions

Next, we assessed lipid droplet deposition using cryosections stained with Sudan III. The specimens included COVID-19–associated pneumonias, non-COVID-19–associated pneumonias and non-pneumonia control lung specimens. The non-pneumonia lungs predominantly were Sudan III-negative (Figure 2A) or contained only single Sudan III-positive cells and extracellular droplets (Figure 2B). Intra-vessel lipid droplets were not detected in non-pneumonia specimens. In the inflamed lung, lipid droplets were visible inside cells, in the alveolar and vessel walls, and inside blood vessels. The pattern was similar for COVID-19–associated (Figure 2C–E) and non-COVID-19–associated pneumonias (Supplementary Figure S5A–C).

**Figure 2.**
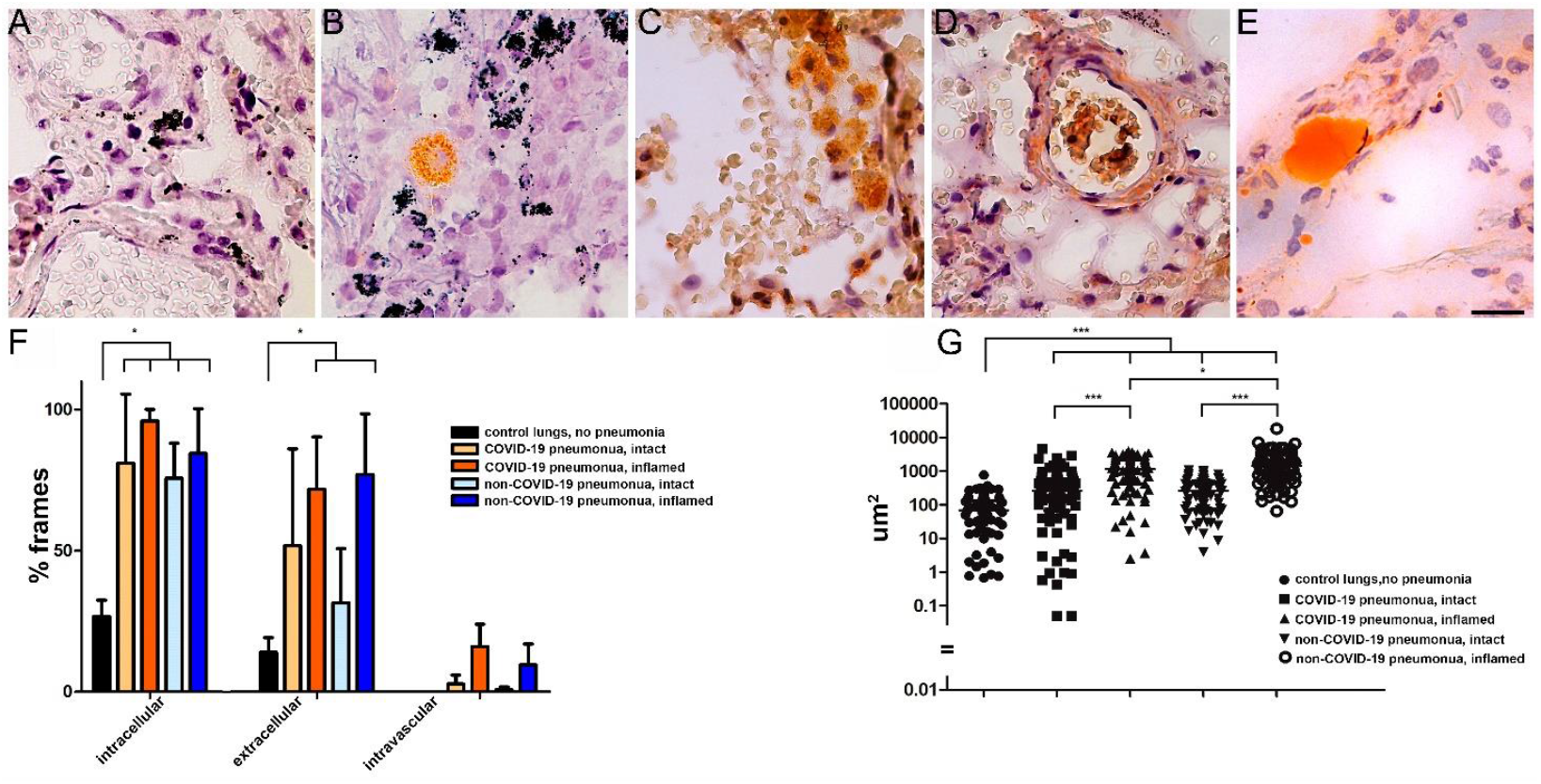
Lipid distribution in non-pneumonia lung tissue and pneumonias of various etiologies. A. Representative non-pneumonia lung tissue staining with Sudan III: no lipid-rich structures are typically present inside blood vessels, in the vessel walls, or in the alveolar intercepts. Scale bar, 25 μm. B. Representative non-pneumonia lung tissue staining with Sudan III: single cells with lipid-rich cytoplasm in the interalveolar wall present the most common type of Sudan III-positive events in non-pneumonia lung tissue. An example of intracellular lipid deposition is shown. Scale bar, 25 μm. C. Representative pneumonia lung tissue staining with Sudan III (COVID-19–associated pneumonia, inflamed area of the lung): grouped cells with lipid-rich cytoplasm are present in the alveolar intercept and in the vicinity of blood vessels. An example of intracellular lipid deposition is shown. Scale bar, 25 μm. D. Representative pneumonia lung tissue staining with Sudan III (COVID-19-associated pneumonia, inflamed area of the lung): the blood vessel wall contains lipid-rich structures among the elastic fibers. An example of extracellular lipid deposition is shown. Scale bar, 25 μm. E. Representative pneumonia lung tissue staining with Sudan III (COVID-19–associated pneumonia, inflamed area of the lung): blood vessel filled with a lipid droplet. An example of intravascular lipid deposition is shown. Scale bar, 25 μm. F. The occurrence rate of intacellular / extracellular / extravascular lipid deposition in non-pneumonia lung tissue (*n*=3 specimens), COVID-19–associated pneumonia (*n*=4 specimens), and non-COVID-19–associated pneumonia (*n*=4 specimens). Analysis performed on 100 consecutive frames per specimen, occurrence rate determined as percentage of frames containing either type of lipid deposition. Inflamed area of the lung (determined macroscopically) was analyzed along with the intact area of the lung (determined macroscopically and obtained from the upper lobe of the same lung) for each pneumonia case. * statistically significant differences (*p*<0.05, Mann-Whitney test). G. The total area of lipid deposition per frame (μm^2^) in non-pneumonia lung tissue (*n*=3 specimens), COVID-19–associated pneumonia (*n*=4 specimens), and non-COVID-19–associated pneumonia (*n*=4 specimens). The inflamed area of the lung was analyzed along with the intact area of the same lung for each pneumonia case. The total area of lipid deposition was assessed on 20 random frames per specimen; the data from specimens were pooled in their respective groups for further analysis. *, statistically significant differences (*p*<0.05, Mann-Whitney test); ***, statistically significant differences (*p*<0.001, Mann-Whitney test).

For quantitative analysis, specimens of non-pneumonia lungs (n=3), COVID-19–associated (n=4), and non-COVID-19–associated (n=4) pneumonia were analyzed. To spatially analyze lipid deposition during inflammation, we assessed two specimens from different sites in the same lung for each pneumonia case: inflamed tissue from the lower lobe and macroscopically intact tissue from the upper lobe. The average occurrence of lipid droplets with different localizations was calculated per 100 consequential high-power fields for each specimen and presented as percentage of fields containing intracellular, extracellular, and intravessel droplets (Supplementary Table S5). The number of cells with cytoplasmic lipid droplets was threefold higher in pneumonia specimens (intact and inflamed lung tissue) than in the control non-pneumonia lung tissue, whereas the number of extracellular lipid droplets was fivefold higher in inflamed pneumonia lung tissue than in the control non-pneumonia lung tissue (Figure 2F), the differences were statistically significant (Mann-Whitney test, p<0.05). Importantly, intra-vessel lipid droplets were detected in intact and inflamed pneumonia lung tissue, but not in the non-pneumonia control. No significant differences (Mann-Whitney test, p>0.05) in the occurrence of lipid droplets were observed for macroscopically intact vs. inflamed tissues from the same lung or for COVID-19–associated vs. non-COVID-19–associated pneumonia (Figure 2F).

We performed a semi-automated morphometric analysis of the total lipid deposition area (for a representative example of the ROIs, see Supplementary Figure S6). No significant differences in total lipid deposition were found between individual specimens within the groups of non-pneumonia lungs, intact pneumonia lungs, and inflamed pneumonia lungs (1-way ANOVA, Dunns post-test). Therefore, for further analysis, data from individuals were pooled into their respective groups (Figure 2G; Supplementary Table S5). The area of lipid deposition was 63.4±20.5 μm^2^ (mean±SD) per frame in the control non-pneumonia lung tissue, 214.5±170.4 μm^2^ in the intact tissue of non-COVID-19– associated pneumonia, 1,732.3±1,212.0 μm^2^ in the inflamed tissue of non-COVID-19–associated pneumonia, 368.5±67.1 μm2 in the intact tissue of COVID-19–associated pneumonia, and 1,494.4±608.6 μm^2^ in the inflamed tissue of non-COVID-19–associated pneumonia. The differences between all groups were statistically significant (Mann-Whitney test, p<0.05) (Figure 2G).

### Fluorescence lipid visualization

To further facilitate morphometric assessment of lipid distribution in the lung tissue, we developed a protocol of triple fluorescence staining of lung cryosections: lipids were stained with Oil Red O that was combined with the intracellular anti-CD31 and DAPI staining (see Materials and Methods for details). Using the triple staining, we confirmed the localization of lipid droplets inside cells, in the alveolar and vessel walls, and inside blood vessels (Figure 3A–C).

**Figure 3.**
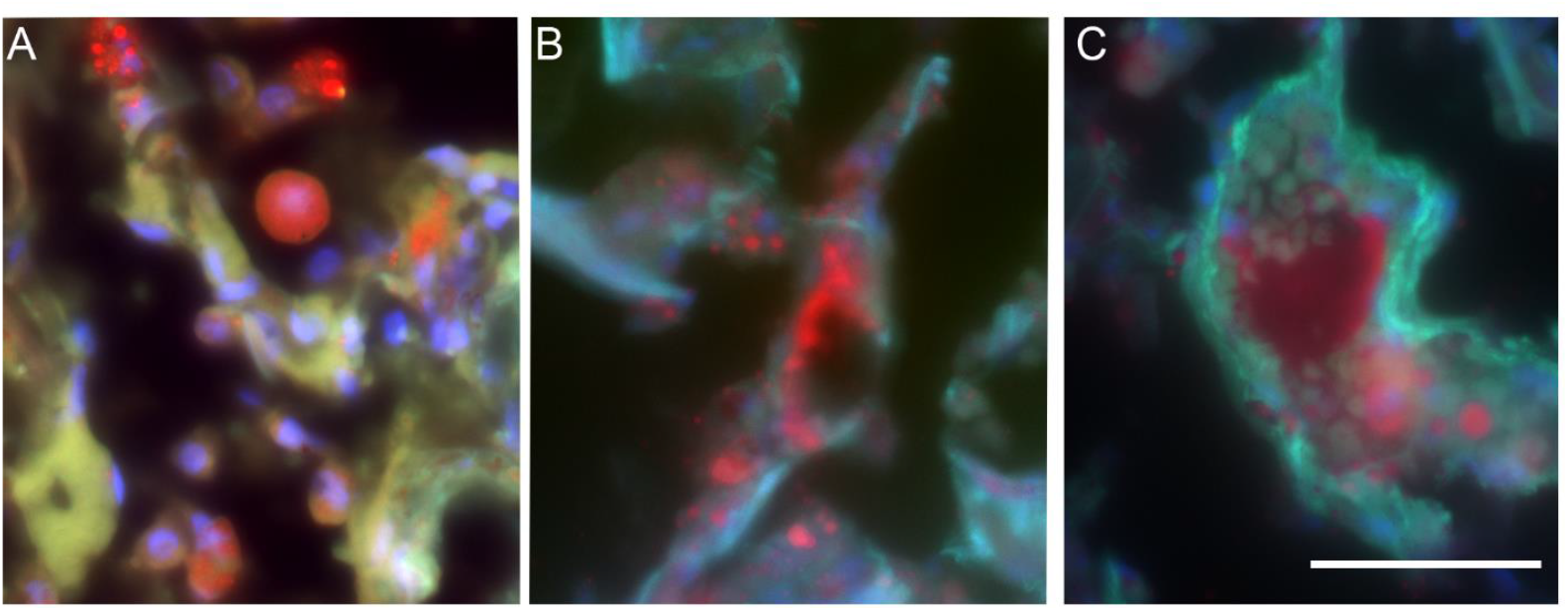
Representative triple fluorescence staining of pneumonia lung tissue with Oil Red O (red)/anti-CD31 (green)/DAPI (blue). To visualize the Oil Red O staining, anti-CD31-FITC, and DAPI staining, the standard Ex.630 nm/Em.670 nm, Ex.488 nm/Em.520 nm, and Ex.405 nm/Em.430 nm filter cubes were used, respectively. Non-COVID-19–associated pneumonia, inflamed area of the lung. Objective x40/Oil. Scale bar, 50 μm. A. Grouped cells with lipid-rich cytoplasm. An example of intracellular lipid deposition is shown. B. Lipid-rich structures in the blood vessel wall. An example of extracellular lipid deposition is shown. Objective x40/Oil. Scale bar, 50 μm. C. blood vessel filled with erythrocytes and a lipid droplet. An example of intravascular lipid deposition is shown. Objective x40/Oil. Scale bar, 50 μm.

### The genes involved in lipid metabolism are differentially expressed in pneumonia lung tissue

Quantitative real-time PCR was used to evaluate the mRNA expression levels of a number of key proteins involved in lipid metabolism and storage in 26 specimens of control and pneumonia lungs, that differed by lipid representation and lipid deposition area. The genes included those coding fatty acid synthase (*FASN)*, lipoprotein lipase (*LPL)*, Long-chain-fatty-acid-CoA ligases (*ACSL1, ACSL3, ACSL4, ACSL5, ACSL6*), phospholipases (*PLA2G4A, PLA2G2A*), perilipins (*PLIN1, PLIN2*). The absence of *PLA2G4A* mRNA expression in the tissues of both control and pneumonia specimen was revealed, therefore this gene was excluded from analysis. The *FASN* gene expression was detected in only two specimens of control lung tissue while in pneumonia lung this mRNA was expressed in the vast majority of specimens. This may indicate the increased expression of *FASN* in pneumonia, however, we were unable to assess the statistical significance for it. The rest of the analyzed mRNAs were detectable both control and pneumonia lung specimens (Figure 4). The difference mRNA expression levels was evaluated using the nonparametric Mann-Whitney test. Three genes showed a statistically significant difference between control and pneumonia lung tissue: *LPL, PLIN2* and *ACSL1*.

**Figure 4.**
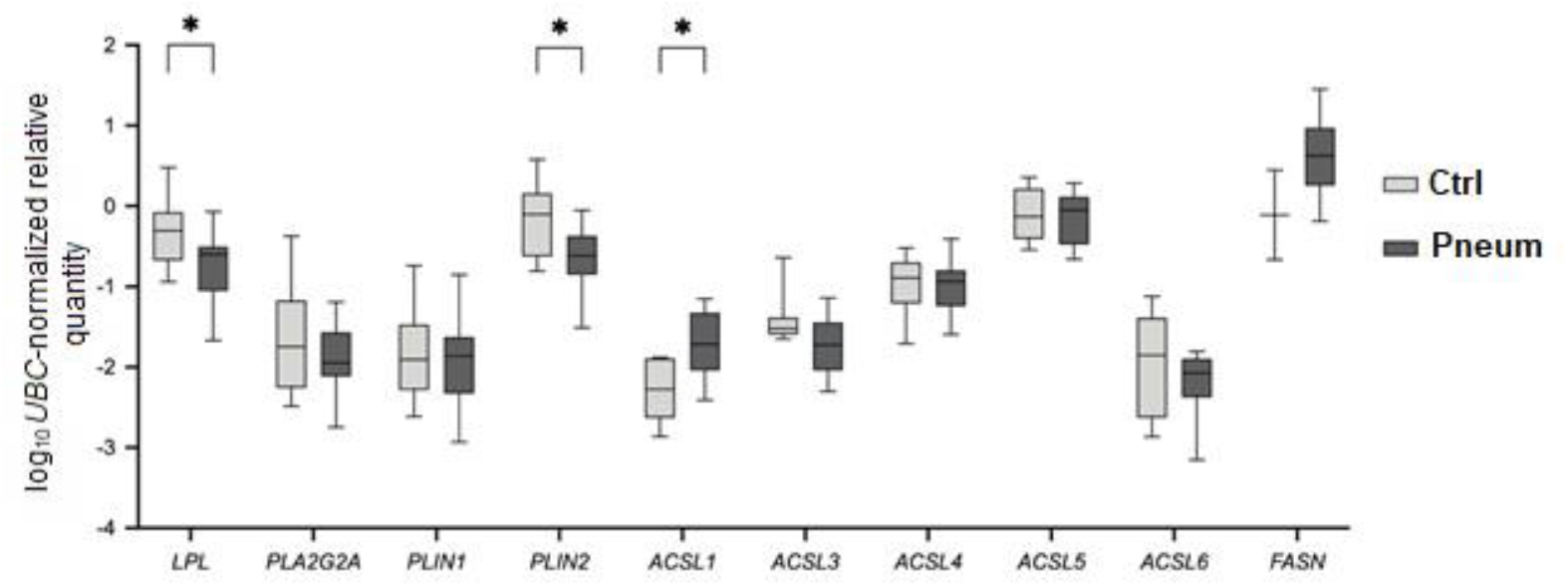
The *UBC*-normalized relative mRNA quantities of genes involved in lipid metabolism as assessed by qPCR. * Statistically significant differences between control (Ctrl) and pneumonia (Pneum) specimens (Mann-Whitney test, p<0.05).

### The FA content allows to distinguish the inflamed vs. the intact tissue from the same lung in pneumonia

As morphometric analysis showed elevated levels of lipid deposition in the inflamed *vs* intact tissue of the same lung in pneumonia we compared the FA content between these sites using chromato-mass spectrometry. While the total FA content was not significantly changed in the inflamed tissue compared with the intact tissue of the same lung (Figure 5A), the unsaturated-to-saturated FA ratio (U/S ratio) was significantly increased (Wilcoxon test, p<0.01) in the specimens with inflammation compared to the intact lung specimens (Figure 5B). Also, we found statistically significant differences (Wilcoxon test, p<0.05) in the content of nine individual FAs in the inflamed vs. the intact tissue from the same lung (Figure 5C). The saturated fatty acids C14:0 (tetradecanoic, myristic), C15:0 (pentadecanoic), C16:0 (palmitic), and C17:0 (heptadecanoic, margaric), and the unsaturated fatty acids C16:1 n-7 (palmitoleic) and C18:3 n-6 (γ-linolenic) were significantly decreased in the inflamed area of the lung compared to the macroscopically intact area, whereas the unsaturated fatty acids C18:1 n-9 (oleic), C18:2 n-6 (α-linoleic), and C22:6 n-3 (docosahexaeoic) were significantly increased in the inflamed area of the lung compared to the macroscopically intact area.

**Figure 5.**
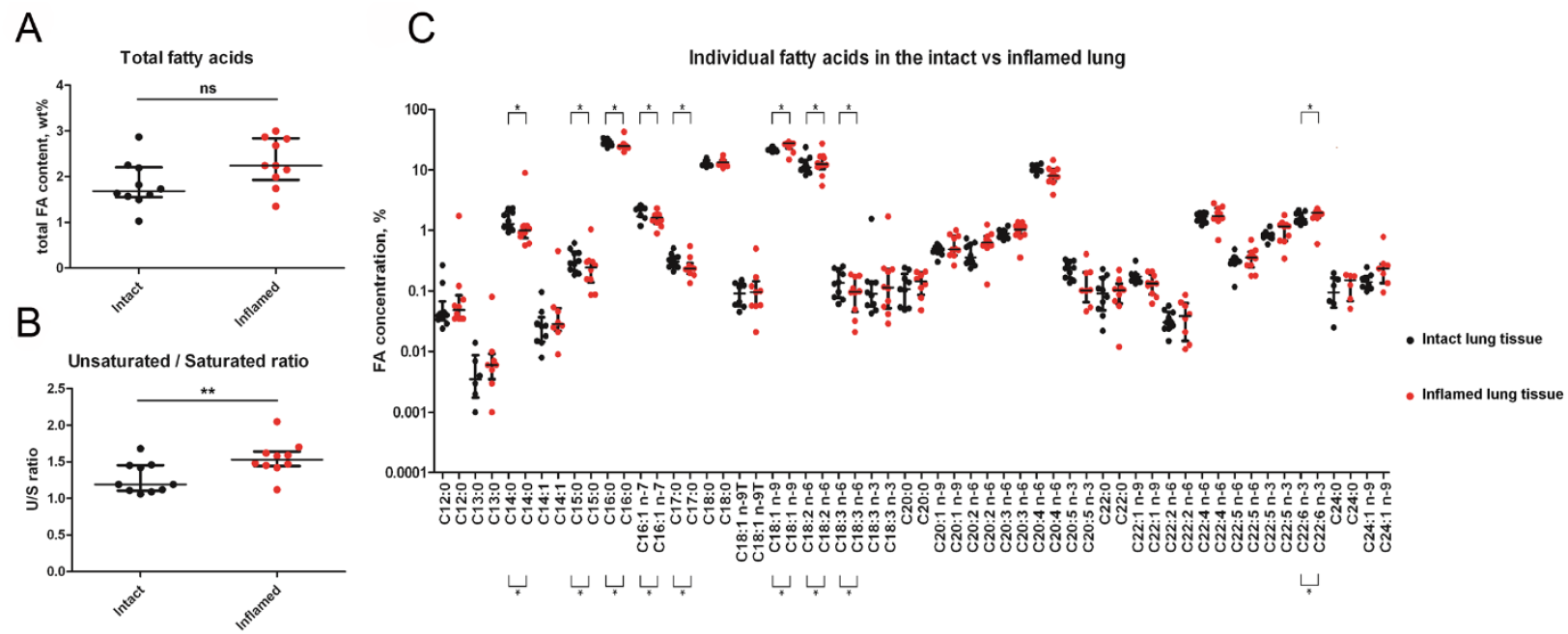
The lipid content of the inflamed vs. intact sections of lung tissue with pneumonia assessed with chromato-mass spectrometry. A. Total fatty acid (FA) content in inflamed pneumonia and intact pneumonia lung specimens. The differences are statistically insignificant. B. The ratio of unsaturated to saturated fatty acids per specimen. The difference is statistically significant (Wilcoxon test, *p*<0.01). C. The concentrations of individual fatty acids. The difference is statistically significant for 9 fatty acids (Wilcoxon test, *p*<0.05). The saturated fatty acids C14:0 (tetradecanoic, myristic), C15:0 (pentadecanoic), C16:0 (hexadecanoic, palmitic), and C17:0 (heptadecanoic, margaric), and the unsaturated fatty acids C16:1 n-7 (palmitoleic) and C18:3 n-6 (γ-linolenic) were significantly decreased in the inflamed area of the lung compared to the macroscopically intact area, whereas the unsaturated fatty acids C18:1 n-9 (oleic), C18:2 n-6 (α-linoleic), and C22:6 n-3 (docosahexaeoic) were significantly elevated in the inflamed area of the lung compared to the macroscopically intact area.

## Discussion

Pathological alterations in pneumonias have been thoroughly studied for over a century. and a number of them have been described [15–17]. The recent COVID-19 pandemic significantly increased pneumonia incidence and triggered a new wave of publications on this disease [18,19]. Nevertheless, among the multiple pathologic processes associated with pneumonia, lipid metabolism remains the least studied. Here, we analyzed the *post mortem* cases with diagnosed COVID-19 pneumonia and non-COVID-19 pneumonia to describe the alterations in lipid deposition in the inflamed lung. Specimens of non-pneumonia lungs served as negative controls.

Lipid depositions were first described morphologically in tuberculosis-associated pneumonia for sites adjacent to the caseous necrosis area in the beginning of the 20th century [20] and were mentioned later in papers describing abnormal lung morphology [21–24]. The scarcity of morphological studies of lipids is partly due to the fact that routine paraffin sections are unsuitable for revealing lipid-rich structures. We first addressed this issue using semi-thin sections with OsO_4_ post-fixation, which stabilizes the lipids [25]. The lipid droplets indeed are best visualized on semi-thin sections stained with methylene blue/azure II. We found small lipid droplets localized within cell cytoplasm and extracellular lipid droplets, most notably large homogeneous intra-vessel lipid droplets. The latter droplets would apparently block the blood flow and affect gas exchange. one of the key aspects of various pneumonias [19, 26], thus significantly contributing to disease progression.

The disadvantage of the semi-thin specimen analysis, however, is that it does not support quantitative analysis due to the small size of the sections. We thus re-addressed this issue using regular histological sections stained with lipophilic dyes and morphometric analysis and showed a significant increase of lipid deposition areas in pneumonias regardless of their etiology. Moreover, we observed a gradient of lipid deposition from the inflamed to the intact region of the same lung. In several studies, the authors also found lipid depositions in COVID-19 pneumonias [23,24] with alternative techniques but related this to systemic factors due to patients’ obesity [22]. In contrast, our data suggest that such abnormalities are local, independent from obesity and the pathogenic factor but characteristic of inflammation in general.

We analyzed the mRNAs of the key regulators of lipid metabolism and storage. Among them were the fatty acid synthase (*FASN*), lipoprotein lipase (*LPL*), long-chain-fatty-acid-CoA ligases (*ACSL1, ACSL3, ACSL4, ACSL5, ACSL6*), phospholipases (*PLA2G4A, PLA2G2A*) and perilipins (*PLIN1, PLIN2*). The expression of *PLIN2* was significantly decreased in the pneumonic lungs compared to control. The PLIN2 protein (also called ADRP) belongs to the perilipin family that binds to the surface of the lipid droplets. The increase in PLIN2 expression leads to the accumulation of lipid droplets and tryglycerides and prevents their hydrolysis [27,28]. Its lack of expression in pneumonia may point to lipid droplet destabilization and causes their extracellular spreading.

The *LPL* mRNA is significantly decreased while *FASN* expression appears to be elevated in pneumonia specimens. Both findings are in accordance with RNA-seq data (Supplementary Table S4) where *FASN* upregulation was attributed to alveolar type II cells and the *LPL* decrease to macrophages. The LPL enzyme catalyzes the cleavage of TGs from chylomicrons and VLDL [29]. FASN is one of the key enzymes in *de novo* fatty acid synthesis [30]. Thus, it can be assumed that fatty acids may be synthesized *de novo* at least by some cell types in the pneumonia lung rather than come from lipoproteins.

The differential fatty acid representation in pneumonia lungs is intriguing and requires a more profound consideration. Apart from specific FA enrichment of the triglycerides upregulated in pneumonia we were able to show that the distribution of several fatty acids follows a gradient in the inflamed lung. Chromato-mass spectrometry revealed a significant shift towards unsaturated FAs in the inflamed lung areas compared to the intact areas of the same lung confirming that lipid abnormalities are local. These data are in agreement with the lipid level changes in bronchoalveolar lavage fluids associated with ARDS and pneumonia, where a decrease in saturated C16:0 and an increase in unsaturated C18:1 and C18:2 have been reported [31,32].

Thus, the morphological and biochemical changes in the inflamed lung both follow a gradient. This gradient may be caused by factors spreading outside the inflamed region. Alternatively, lipid metabolism abnormalities may constitute an early event in lung inflammation and tissue in the upper lung segments, while still macroscopically intact, present the first pneumonia-caused changes. The changes in lipid deposition may be caused by inflammation-driven shifts in the expression of genes involved in lipid synthesis and turnover [33,34]. In addition. lipid accumulation may result from tissue decomposition in the inflamed areas, the emergence and the extracellular spread of lipid droplets combined with the macrophages’ inability to eliminate the accumulated lipids [35,36].

In conclusion, our data provide new insights into pneumonia pathogenesis and its association with lipid metabolism and inflammation. Our results indicate that lipid metabolism can be considered a target for therapeutic interventional strategy in pneumonia.

