## Supplementary Tables and Figures for "Lung lipid deposition in pneumonias of different etiologies"

### Supplementary Figures

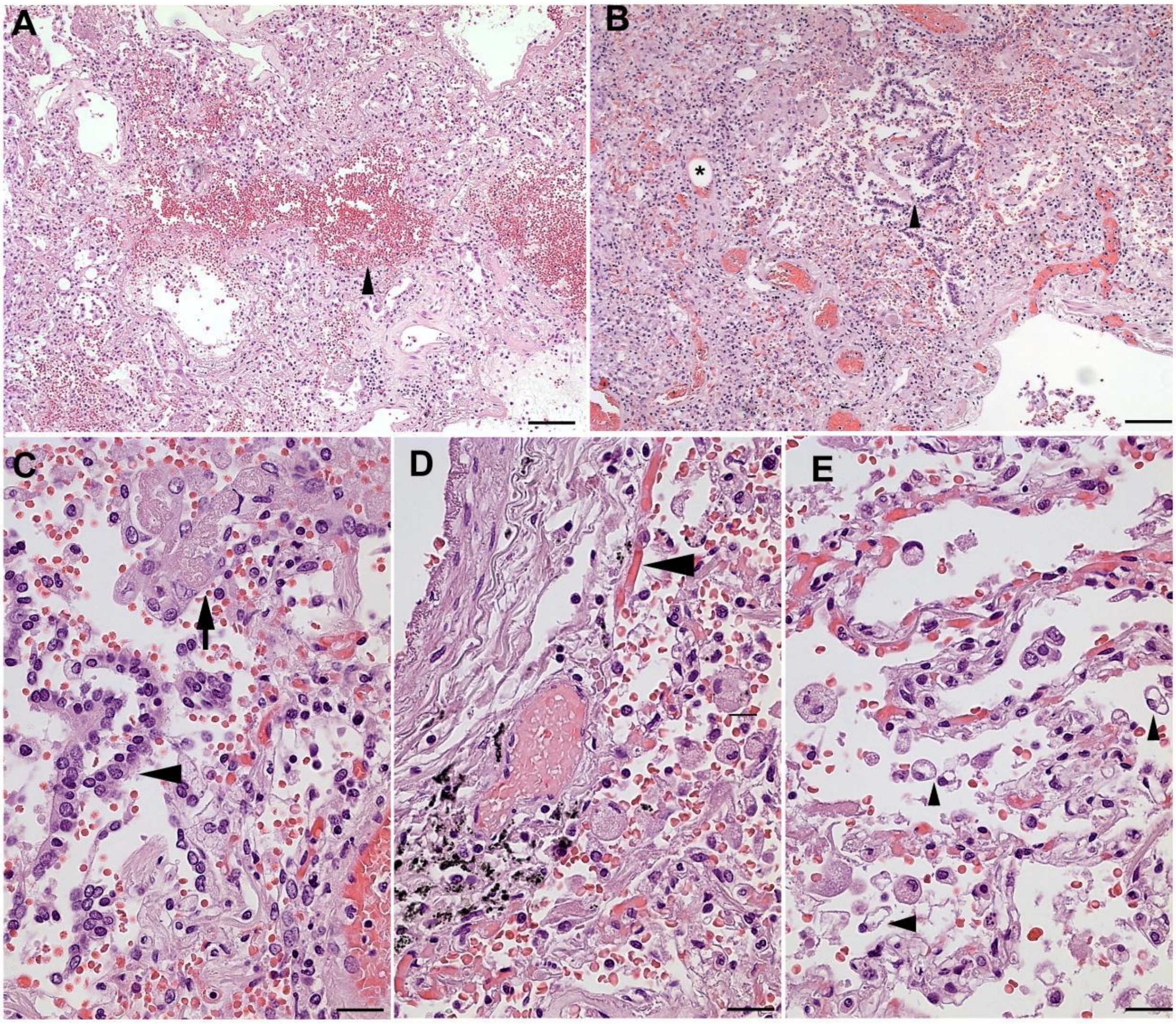

**Supplementary Figure S1** H&E paraffin sections of COVID-19–associated pneumonias, characteristic changes of lung tissue morphology. A. Exudative stage of diffuse alveolar damage, loss of lung morphology; fields of extravascular erythrocytes within lung tissue (arrowhead). Scale bar, 100 μm. B. Proliferative stage of diffuse alveolar damage, loss of lung morphology; alveolar remnants contain desquamed epithelial cells (arrowhead). The blood vessels are filled with erythrocytes; surrounded by erythrocytes is the cavity that presumably contained a lipid droplet (asterisk). Scale bar, 100 μm. C. Alveolar remnants with giant cells (arrow) and desquamed epithelium (arrowhead). Scale bar, 25 μm. D. Blood vessels filled with erythrocytes. Characteristic erythrocyte sludges in capillaries (arrowhead). Scale bar, 25 μm. E. Cells with foamed cytoplasm or empty cavities in the cytoplasm (arrowheads). Scale bar, 25 μm

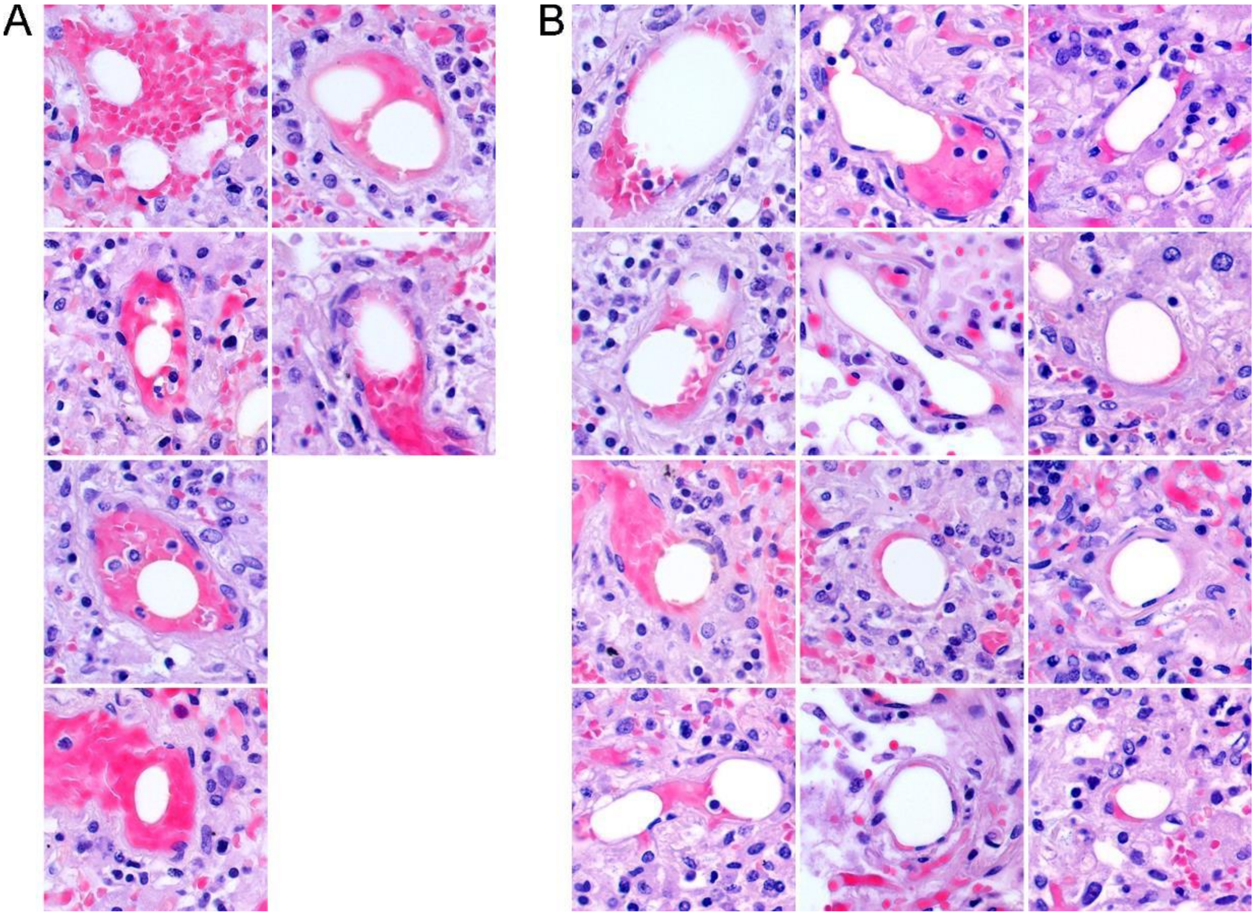

**Supplementary Figure S2** The image gallery obtained upon the examination of a single random H&E specimen of COVID-19–associated pneumonia. The majority of the blood vessels in pneumonia lungs are filled with erythrocytes. A. A large number of vessels contain cavities, completely surrounded by erythrocytes, which presumably contained lipid-rich structures lost upon specimen dehydration. B. Other vessels contain deformed erythrocytes pushed to the side of the vessel. The sizes of the cavities are close to the sizes of small vessels.

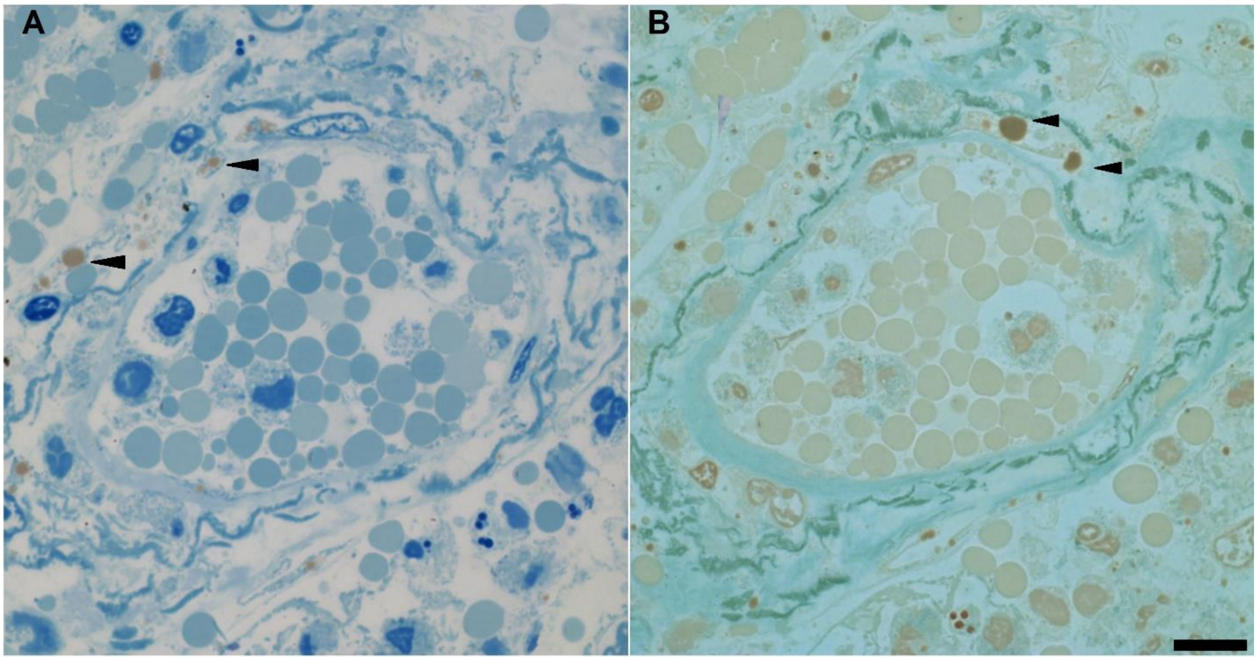

**Supplementary Figure S3** Comparison of methylene blue/azure II and Twort's staining of lipid-rich structures in COVID-19–associated pneumonia. A. The lipid depositions are clearly visible in lung tissue methylene blue/azure II specimens; the lipid-rich structures in this case have brown color (arrowheads) that distinguishes them from other cellular and extracellular structures, which have blue color. B. The color differentiation between lipid droplets and erythrocytes is not so obvious with Twort's staining: the lipid droplets in this case have dark brown color (arrowheads), while the erythrocytes have pale brown color. Scale bar, 10 µm.

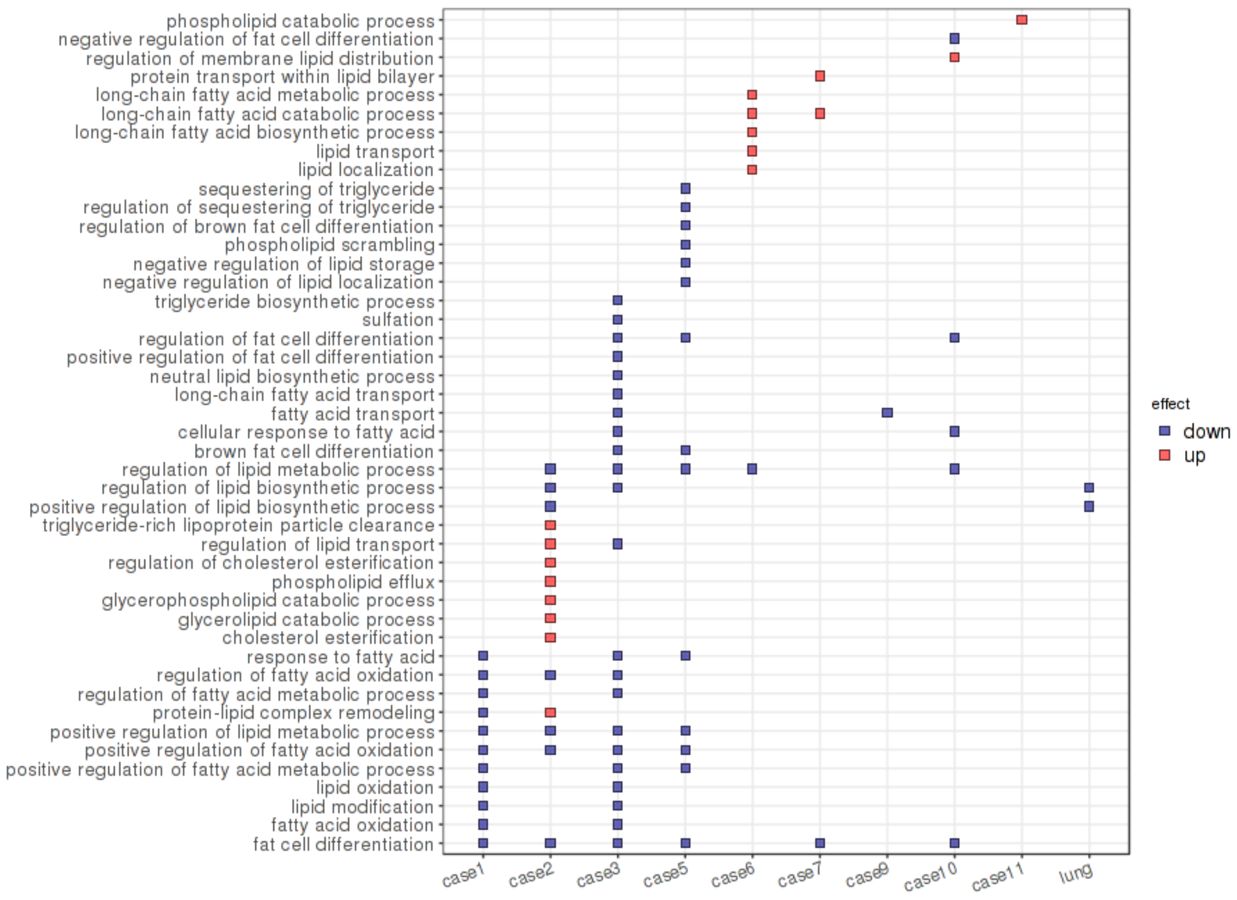

**Supplementary Figure S4** GO analysis: the bulk RNA-seq dataset included 9 independent pneumonia cases ("case1"; "case2"; "case3"; "case5"; "case6"; "case7"; "case9"; "case10"; "case11") and the data on the cumulative dataset ("lung"). The alterations were found in a number of gene clusters, including those specified by GO terms "regulation of lipid biosynthetic process" and "positive regulation of lipid biosynthetic process" (for the cumulative dataset), "fat cell differentiation" (for 6 of 9 cases), and "regulation of lipid metabolic process" (for 5 of 9 cases). The overall trend for gene downregulation is observed despite the variability between cases.

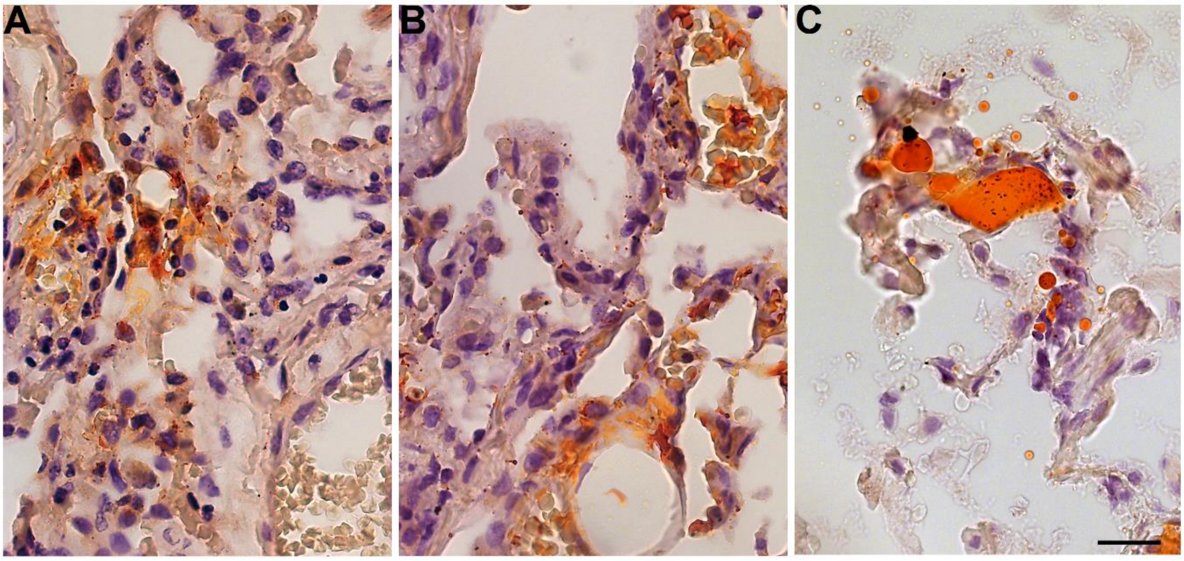

**Supplementary Figure S5** The lipid distribution in non-COVID-19–associated pneumonia. A. Representative pneumonia lung tissue staining with Sudan III (non-COVID-19–associated pneumonia, inflamed area of the lung): grouped cells with lipid-rich cytoplasm are present in the alveolar intercept and in the vicinity of blood vessels. An example of intracellular lipid deposition is shown. Scale bar, 25 µm. B. Representative pneumonia lung tissue staining with Sudan III (non-COVID-19-associated pneumonia, inflamed area of the lung): the blood vessel wall contains lipid-rich structures among the elastic fibers. An example of extracellular lipid deposition is shown. Scale bar, 25 µm. C. Representative pneumonia lung tissue staining with Sudan III (non-COVID-19–associated pneumonia, inflamed area of the lung): blood vessel filled with a lipid droplet. An example of intravascular lipid deposition is shown. Scale bar, 25 µm.

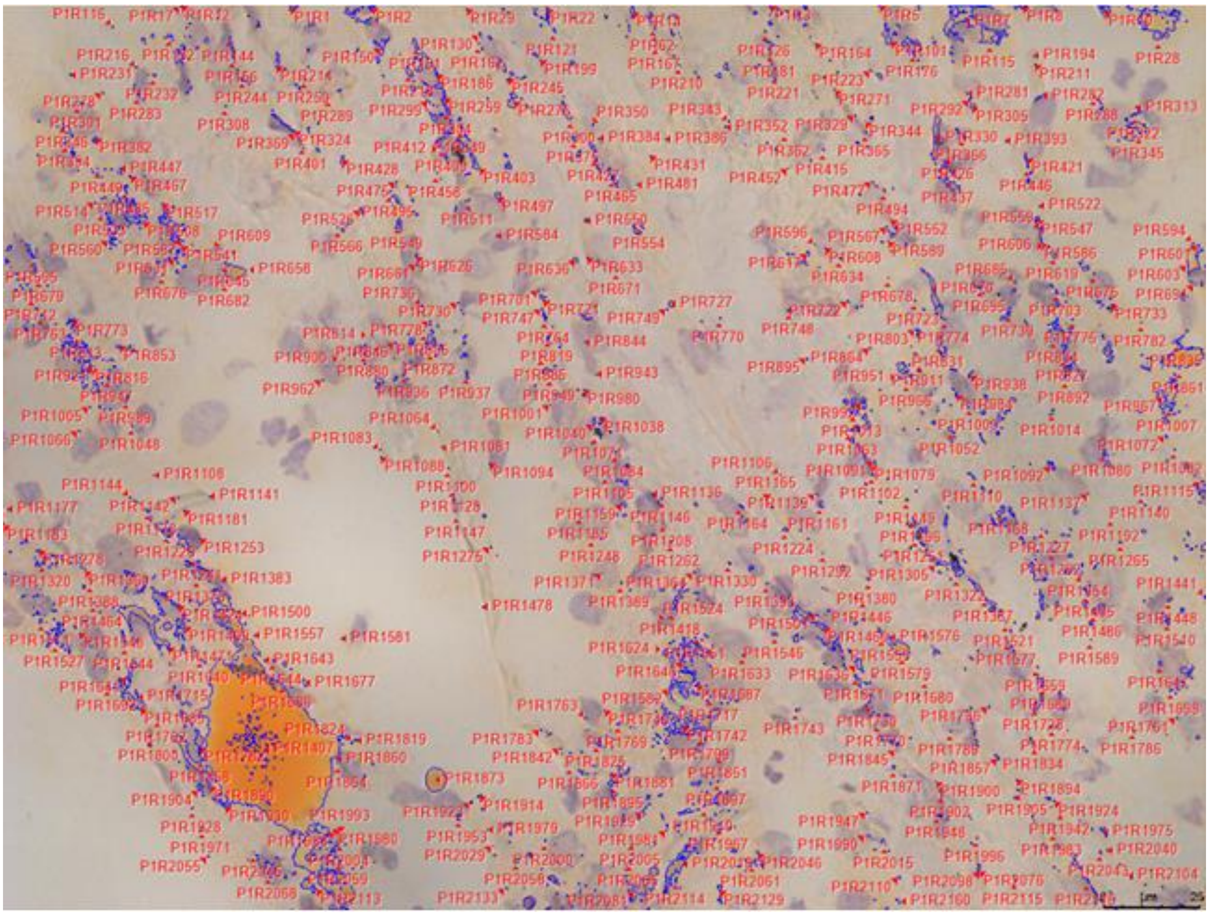

**Supplementary Figure S6** Example of the morphometric analysis of Sudan III-stained lung cryosections. For each image, Sudan III–positive droplets were included in an ImagePro mask containing pixels by color. The mask was created semi-automatically based on the operator-chosen pixel color, and manually adjusted to include all Sudan III–positive structures into the regions of interest (ROIs). The total area of the mask was calculated in μm². Scale bar, 25 μm

**Supplementary Tables**

**Supplementary Table S1.** Description of the autopsies included in the study.

| Autopsy | Sex | Age | Pneumonia | COVID-19 in-hospital qPCR | COVID-19 post-mortem qPCR | BMI |
|---|---|---|---|---|---|---|
| **1 COVID-19** | F | 82 | + | NA | + | BMI<30 |
| **2 COVID-19** | M | 79 | + | + | + | NA |
| **3 COVID-19** | F | 68 | + | NA | + | BMI≥30 |
| **4 COVID-19** | F | 61 | + | + | + | BMI≥30 |
| **5 COVID-19** | M | 79 | + | + | + | NA |
| **6 COVID-19** | M | 62 | + | + | + | NA |
| **7 COVID-19** | M | 73 | + | + | + | BMI≥30 |
| **8 COVID-19** | M | 79 | + | + | + | BMI≥30 |
| **9 COVID-19** | F | 78 | + | - | + | BMI<30 |
| **10 COVID-19** | M | 73 | + | - | + | BMI<30 |
| **11 COVID-19** | M | 89 | + | + | + | BMI≥30 |
| **12 COVID-19** | M | 74 | + | + | + | BMI<30 |
| **13 COVID-19** | F | 84 | + | + | + | BMI<30 |
| **14 COVID-19** | F | 70 | + | + | NA | BMI<30 |
| **15 COVID-19** | M | 84 | + | + | NA | BMI<30 |
| **16 COVID-19** | F | 67 | + | + | NA | 24,7 |
| **17 COVID-19** | M | 62 | + | - | + | BMI<30 |
| **18 COVID-19** | M | 61 | + | - | + | 15,6 |
| **19 COVID-19** | F | 87 | + | NA | + | BMI<30 |
| **20 COVID-19** | M | 71 | + | - | + | BMI<30 |
| **21 COVID-19** | M | 71 | + | - | + | NA |
| **22 COVID-19** | F | 63 | + | - | + | 31,5 |
| **23 COVID-19** | F | 80 | + | NA | + | 18.5 |

| | | | | | | |
|---|---|---|---|---|---|---|
| **1 non-COVID-19** | M | 54 | + | - | - | 24,8 |
| **2 non-COVID-19** | M | 67 | + | - | - | BMI<30 |
| **3 non-COVID-19** | M | 40 | + | - | - | 32,8 |
| **4 non-COVID-19** | F | 76 | + | - | - | BMI<30 |
| **5 non-COVID-19** | F | 47 | + | - | - | 28 |
| **6 non-COVID-19** | M | 61 | + | - | - | BMI<30 |
| **7 non-COVID-19** | M | 89 | + | - | - | BMI<30 |
| **8 non-COVID-19** | F | 89 | + | - | - | 19,5 |
| **9 non-COVID-19** | F | 73 | + | - | - | 28 |
| **10 non-COVID-19** | M | 89 | + | - | - | 22 |
| **11 non-COVID-19** | M | 75 | + | - | - | 23,1 |
| **12 non-COVID-19** | M | 82 | + | - | - | BMI<30 |
| **13 non-COVID-19** | M | 81 | + | - | - | 24,6 |
| **14 non-COVID-19** | F | 59 | + | - | - | 18.5 |
| **15 non-COVID-19** | F | 80 | + | NA | - | 33,7 |
| **16 non-COVID-19** | F | 62 | + | NA | - | 37,2 |
| **17 non-COVID-19** | F | 61 | + | - | - | 28,4 |
| **18 non-COVID-19** | M | 68 | + | NA | - | 25,3 |
| **1 Ctrl** | F | 95 | - | NA | - | BMI<30 |
| **2 Ctrl** | M | 90 | - | NA | - | BMI<30 |
| **3 Ctrl** | F | 74 | - | NA | - | BMI<30 |
| **4 Ctrl** | F | 67 | - | NA | - | BMI<30 |
| **5 Ctrl** | M | 57 | - | NA | - | BMI<30 |
| **6 Ctrl** | F | 64 | - | NA | - | BMI<30 |
| **7 Ctrl** | F | 84 | - | NA | - | BMI<30 |
| **8 Ctrl** | M | 84 | - | NA | - | 32,4 |
| **9 Ctrl** | F | 73 | - | NA | - | 20,2 |
| **10 Ctrl** | M | 69 | - | NA | - | 32,1 |

*BMI, body mass index. NA, data not available.

**Supplementary Table S2.** qPCR primers and probes for SARS-CoV-2 detection in lung tissue.

| Name | Sequence | Label |
|---|---|---|
| ***N2*-forward** | TTACAAACATTGGCCGCAAA | - |
| ***N2*-reverse** | GCGCGACATTCCGAAGAA | - |
| ***N2*-probe** | ACAATTTGCCCCCAGCGCTTCAG | 5’-FAM 3’-BHQ1 |
| ***N3*-forward** | GGGAGCCTTGAATACACCAAAA | - |
| ***N3*-reverse** | TGTAGCACGATTGCAGCATTG | - |
| ***N3*-probe** | ATCACATTGGCACCCGCAATCCTG | 5’-VIC 3’-BHQ2 |
| ***UBC*-forward** | TTGGGTCGCAGTTCTTGTTTG | - |
| ***UBC*-reverse** | TGCCTTGACATTCTCGATGGT | - |
| ***UBC*-probe** | TCGCTGTGATCGTCACTTGACAATG | 5’-ROX 3’-BHQ2 |

*The *N2* and *N3* sequences used were recommended for SARS-CoV-2 detection by the Division of Viral Diseases, National Center for Immunization and Respiratory Diseases, Centers for Disease Control and Prevention, Atlanta, GA, USA. *UBC* was added as a reference gene. All oligonucleotides were synthesized with DNK-Sintez (Russia) and put into a triplex one-step qPCR reaction. Sequences are presented in 5′–3′direction. FAM, fluorescein; VIC, 2′-chloro-7′phenyl-1,4-dichloro-6-carboxy-fluorescein; ROX, carboxyrhodamine; BHQ1/2, Black Hole Quencher 1/2.

**Supplementary Table S3.** qPCR primers for the SYBR Green I detection of genes of interest in lung tissue.

| Name | Sequence |
|---|---|
| ***COX1*-forward** | TTCTTGCTGTTCCTGCTCCT |
| ***COX1*-reverse** | TGACAAACTCCCAGAACCAG |
| ***COX2*-forward** | GATGATTGCCCGACTCCCTT |
| ***COX2*-reverse** | CTAGCCAGAGTTTCACCGTAA |
| ***LPL*-forward** | GTGCCAGAACTTCGACCCTT |
| ***LPL*-reverse** | TTTTCGGAGCCAGAGTTCCC |
| ***PLA2G4A*-forward** | ATCACACGAACCCAAAGGCA |
| ***PLA2G4A* -reverse** | AGCCCAGCATGAAGTTGTGT |
| ***PLA2G2A*-forward** | GGAGTGCAGAACAAACAAGAC |
| ***PLA2G2A* -reverse** | GCAGCCGTAGAAGCCATAAC |
| ***ACSL1-forward*** | CTGGTACGCCACGAGACC |
| ***ACSL1-reverse*** | TCGTCGCTGTCAAGTAGTGC |
| ***ACSL3-forward*** | TGGATGATAGCTGCACAGGC |
| ***ACSL3-reverse*** | GCGTGGGACCAAAGAAACTATA |
| ***ACSL4-forward*** | TGTGAAAGAATACCTGGACTGG |
| ***ACSL4-reverse*** | GCCATAAGTGTGGGCTTCAG |
| ***ACSL5-forward*** | CAAAGAGGACTCGCTGTGTC |
| ***ACSL5-reverse*** | GTAACAAGCCAATTCGGAGATG |
| ***ACSL6-forward*** | GCTTCACACTGGAGACATCG |
| ***ACSL6-reverse*** | GAGGGCATAACTTCAGGGTC |
| ***FASN-forward*** | TCCTTCTTCGGAGTCCACC |
| ***FASN-reverse*** | TCCTCGGAGTGAATCTGGGT |
| ***PLIN1-forward*** | CCTGCTGTCTCCTCAACCAA |
| ***PLIN1-reverse*** | GCGTGGCGACTTCGTCCTC |
| ***PLIN2-forward*** | ACAACCGAGTGTGGTGACTC |
| ***PLIN2-reverse*** | TCCTGTCTAGCCCCTTACAG |
| ***UBC*-forward** | TTGGGTCGCAGTTCTTGTTTG |

| | |
|---|---|
| ***UBC*-reverse** | TGCCTTGACATTCTCGATGGT |

* All oligonucleotides were synthesized by DNK-Sintez (Russia) and used in qPCR reactions to analyze cDNA obtained from lung tissue. Sequences are presented in 5′–3′ direction.

**Supplementary Table S4**. Single-cell RNAseq analysis.

| Cell Type | Tissue | Gene ID | Up/Down |
|---|---|---|---|
| **Alveolartype II cells** | Lung | *MSMO1* | Upregulated |
| **Alveolartype II cells** | Lung | *SCD* | Upregulated |
| **Alveolartype II cells** | Lung | *FDFT1* | Upregulated |
| **Alveolartype II cells** | Lung | *HMGCS1* | Upregulated |
| **Alveolartype II cells** | Lung | *FASN* | Upregulated |
| **Fibroblasts** | Lung | *FGF2* | Downregulated |
| **Fibroblasts** | Lung | *RARRES2* | Downregulated |
| **Fibroblasts** | Lung | *DKK3* | Downregulated |
| **Fibroblasts** | Lung | *PDGFRA* | Downregulated |
| **Fibroblasts** | Lung | *FGFR4* | Downregulated |
| **Endothelialcells** | Lung | *FLT1* | Upregulated |
| **Endothelialcells** | Lung | *RELN* | Upregulated |
| **Endothelialcells** | Lung | *PIK3R3* | Downregulated |
| **Endothelialcells** | Lung | *PDGFB* | Downregulated |
| **Endothelialcells** | Lung | *FGF2* | Downregulated |
| **Macrophages** | Lung | *PSAP* | Upregulated |
| **Macrophages** | Lung | *SPP1* | Upregulated |
| **Macrophages** | Lung | *GLUL* | Upregulated |
| **Macrophages** | Lung | *FABP4* | Downregulated |
| **Macrophages** | Lung | *RBP4* | Downregulated |
| **Macrophages** | Lung | *RETN* | Downregulated |
| **Macrophages** | Lung | *INHBA* | Downregulated |
| **Macrophages** | Lung | *PPARG* | Downregulated |
| **Macrophages** | Lung | *FFAR4* | Downregulated |
| **Macrophages** | Lung | *LPL* | Downregulated |
| **Macrophages** | Lung | *FABP3* | Downregulated |

| | | | |
|---|---|---|---|
| **Macrophages** | Lung | *NMB* | Downregulated |
| **Macrophages** | Lung | *ADTRP* | Downregulated |
| **Macrophages** | Lung | *AKR1C2* | Downregulated |
| **Macrophages** | Lung | *SCD* | Downregulated |
| **Macrophages** | Lung | *AKR1C3* | Downregulated |
| **Monocytes** | Lung | *ABHD5* | Upregulated |
| **Monocytes** | Lung | *G0S2* | Upregulated |
| **Monocytes** | Lung | *HCAR2* | Upregulated |
| **Monocytes** | Lung | *ANXA2* | Downregulated |
| **Monocytes** | Lung | *THBS1* | Downregulated |
| **T cells** | Lung | *IFNG* | Upregulated |
| **T cells** | Lung | *FYN* | Upregulated |
| **T cells** | Lung | *CCL5* | Upregulated |
| **T cells** | Lung | *LIPA* | Downregulated |
| **T cells** | Lung | *FUCA1* | Downregulated |
| **T cells** | Lung | *APOE* | Downregulated |
| **T cells** | Lung | *APOC1* | Downregulated |
| **T cells** | Lung | *APOC2* | Downregulated |
| **T cells** | Lung | *PLTP* | Downregulated |
| **T cells** | Lung | *FABP3* | Downregulated |
| **T cells** | Lung | *FABP4* | Downregulated |
| **T cells** | Lung | *SCD* | Downregulated |

* The tissue type in analysis was restricted to “Lung” and the cell types in analysis were restricted to the most prominent resident/infiltrating cell populations. The genes showing controversial results in different datasets were excluded. In the lung tissue, the downregulated genes in lung tissue are mostly assigned to fibroblasts (5 downregulated genes), macrophages (13 downregulated genes, 3 upregulated genes), and T cells (9 downregulated genes, 3 upregulated genes).

**Supplementary Table S5.** Lipid droplet distribution in lung tissue; morphologic and morphometric analysis.

| Specimen | Number of frames with: | | | Total area of lipid droplets, μm$^2$ | |
|---|---|---|---|---|---|
| | Intracellular lipid droplets / per 100 frames | Extracellular lipid droplets / per 100 frames | Intra-vessel lipid droplets / per 100 frames | Median / per 20 high-power fields | IQR* / per 20 high-power fields |
| **Non-pneumonia lung 1** | 30 | 11 | 0 | 39.71 | 10.26-89.42 |
| **Non-pneumonia lung 2** | 20 | 11 | 0 | 76.57 | 27.53-131.33 |
| **Non-pneumonia lung 3** | 30 | 20 | 0 | 73.87 | 31.45-168.94 |
| **Non-COVID-19 pneumonia, intact lung 1** | 65 | 30 | 1 | 327.60 | 236.26-428.28 |
| **Non-COVID-19 pneumonia, intact lung 2** | 93 | 12 | 0 | 75.95 | 51.76-270.78 |
| **Non-COVID-19 pneumonia, intact lung 3** | 76 | 58 | 0 | 404.75 | 199.51-732.86 |
| **Non-COVID-19 pneumonia, intact lung 4** | 69 | 26 | 2 | 162.87 | 74.32-415.88 |
| **Non-COVID-19 pneumonia, inflamed lung 1** | 96 | 89 | 0 | 1,178.58 | 706.10-1672.91 |
| **Non-COVID-19 pneumonia, inflamed lung 2** | 100 | 100 | 18 | 2,930.49 | 2081.97-4,549.38 |
| **Non-COVID-19 pneumonia, inflamed lung 3** | 74 | 67 | 11 | 1,759.32 | 642.48-2,347.01 |
| **Non-COVID-19 pneumonia, inflamed lung 4** | 68 | 52 | 9 | 506.99 | 313.01-1,559.79 |
| **COVID-19 pneumonia, intact lung 1** | 45 | 19 | 1 | 47.97 | 2.74-194.60 |
| **COVID-19 pneumonia, intact lung 2** | 90 | 47 | 7 | 348.75 | 163.46-704.57 |
| **COVID-19 pneumonia, intact lung 3** | 89 | 41 | 3 | 313.58 | 123.13-773.69 |
| **COVID-19 pneumonia, intact lung 4** | 100 | 100 | 0 | 443.24 | 261.63-660.08 |

| | | | | | |
|---|---|---|---|---|---|
| **COVID-19 pneumonia, inflamed lung 1** | 99 | 59 | 5 | 546.49 | 107.84-1,082.08 |
| **COVID-19 pneumonia, inflamed lung 2** | 93 | 72 | 18 | 1,236.84 | 619.31-2,516.79 |
| **COVID-19 pneumonia, inflamed lung 3** | 92 | 58 | 17 | 1,056.92 | 235.39-1,428.66 |
| **COVID-19 pneumonia, inflamed lung 4** | 100 | 98 | 24 | 2,189.44 | 1415.08 – 2,738.02 |

* IRQ, interquartile range
